# Genetic diversity in vector populations influences the transmission efficiency of an important plant virus

**DOI:** 10.1101/2024.02.21.581289

**Authors:** Daniel J Leybourne, Mark A Whitehead, Torsten Will

## Abstract

The transmission efficiency of aphid-vectored plant viruses can differ between aphid populations. Intra-species diversity (genetic variation, endosymbionts) is a key determinant of aphid phenotype; however, the extent of which intra-species diversity contributes towards variation in virus transmission efficiency is unclear. Here, we use multiple populations of two key aphid species that vector barley yellow dwarf virus (BYDV) strain PAV (BYDV-PAV), the grain aphid (*Sitobion avenae*) and the bird cherry-oat aphid (*Rhopalosiphum padi*), and examine how diversity in vector populations influences virus transmission efficiency. We use Illumina sequencing to characterise genetic and endosymbiont variation in multiple *S. avenae* and *R. padi* populations and conduct BYDV-PAV transmission experiments to identify links between intra-species diversity in the vector and virus transmission efficiency. We observe limited variation in the transmission efficiency of *S. avenae*, with transmission efficiency consistently low for this species. However, for *R. padi* we observe a range of transmission efficiency and show that BYDV transmission efficiency is influenced by genetic diversity within the vector, identifying 542 SNPs that potentially contribute towards variable transmission efficiency in *R. padi*. Our results represent an important advancement in our understanding of the relationship between genetic diversity, vector-virus interactions, and virus transmission efficiency.

## Introduction

Cereal aphids, including the grain aphid, *Sitobion avenae*, and the bird cherry-oat aphid, *Rhopalosiphum padi*, are important herbivorous insects of cereal crops [1]. Cereal aphids are widely distributed across Central Europe and cause significant crop damage through feeding [2] and the transmission of plant viruses [3]. Cereal aphids vector several plant viruses, including those that cause yellow dwarf disease [4]. Yellow dwarf disease is caused by multiple viruses, including barley yellow dwarf virus (BYDV, Tombusviridae : *Luteovirus*), cereal yellow dwarf virus (CYDV, Solemoviridae : *Polerovirus*), maize yellow dwarf virus (MYDV, Solemoviridae : *Polerovirus*), and wheat yellow dwarf virus (WYDV, Solemoviridae) [5]. There are several virus species within each yellow dwarf virus genus [4, 5], however the most dominant and agriculturally important in the UK and Europe is BYDV-PAV (Tombusviridae : *Luteovirus pavhordei*) [6]. Infection with BYDV-PAV can decrease crop yield by *c*. 20% [3, 7]. Yellow dwarf disease symptoms include crop stunting, delayed crop maturity, shrivelled grain, reduced transpiration, and chlorosis [5].

Aphid (vector) and disease management strategies for yellow dwarf disease follow strict thresholds [8]. In the UK, the current threshold, the level of aphid infestation above which treatment is recommended, is the presence of a single virus-vectoring aphid (*R. padi, S. avenae*, or the rose-grain aphid *Metapolophium dirhodum*) in the crop during the early stages of plant growth [8, 9]. Once the crop reaches growth stage 31 it is able to naturally tolerate yellow dwarf virus infection [10]. Similar stringent thresholds are followed in other European countries. These low thresholds have likely contributed to increased application of management interventions such as insecticide treatments, directly increasing the development of insecticide resistant, or desensitised, aphid populations [11-13]. Currently, the same yellow dwarf virus threshold applies to all vector species, and all populations within a vector species. This is an important oversight, as aphid populations are not homogenous and there is inherent diversity within vector populations that can significantly influence the behaviour and phenology of both the aphid and the virus. Indeed, the transmission efficiency of BYDV-PAV differs between cereal aphid species, and a recent review found that transmission efficiency can also vary between aphid populations within a given species [5]. For example, transmission efficiency of BYDV-PAV by *R. padi* can range from 39-80%, 0-100%, and 20-100% for wheat, barley, and oats, respectively [5]. The biological drivers behind this variation are poorly understood, however intra-species diversity (aphid genetic diversity and the presence and diversity of aphid endosymbionts) within aphid populations are some proposed hypotheses [5].

The majority of aphid species form an obligatory relationship with the endosymbiont *Buchnera aphidicola*. In this endosymbiotic relationship *B. aphidicola* supplements the diet of the host aphid through provision of amino acids [14]. Diversity within *B. aphidicola* can also influence other aspects of aphid fitness and behaviour, with different *B. aphidicola* strains conferring additional beneficial traits to the aphid host, such as heat tolerance [15]. Aphids can also form a range of non-essential, or facultative, relationships with several endosymbionts [16, 17] that also influence aphid phenotype [18]. The facultative endosymbionts described to associate with aphids include, *Regiella insecticola, Hamiltonella defensa, Fukatsuia symbiotica* (previously PAXS), *Serratia symbiotica, Rickettsia* spp., *Ricketsiella* spp., *Spiroplasma* spp., and *Arsenophonus* spp. [11, 16, 19, 20]. Facultative endosymbionts occur naturally in cereal aphid populations [11, 20] and facultative endosymbiont infections can exist in individual infections, co-infections, or multi-infections [11, 20-22]. Several fitness and behavioural traits can be conferred to the host aphid by facultative endosymbionts, including protection against parasitism by [21] and differential feeding behaviour [23]. Endosymbiont effects can also be mediated by aphid genotype, through an endosymbiont x genotype interaction [22], and aphid genotype inherently influences aphid fitness [21].

Despite the broad effects intra-species diversity (genotype, *B. aphidicola* strain, facultative endosymbiont presence and strain) has on aphid phenology and behaviour relatively few studies have examined how these traits impact aphid-virus interactions. Some recent studies have started to explore the potential influence facultative endosymbionts might have on the aphid-BYDV relationship [24-26], however transmission efficiency is often not directly examined [25] or the observed endosymbiont effects cannot be disentangled from the confounding effect of aphid genotype [24]. Genetic variation has been found to underpin transmission efficiency in another aphid-yellow dwarf virus combination [27], but this remains understudied for *R. padi, S. avenae*, and BYDV-PAV. Here, we use Illumina sequencing to characterise genetic and endosymbiont diversity in aphid populations and combine this with BYDV transmission experiments to examine how diversity in vector populations impacts BYDV transmission efficiency. To achieve this, we use the most prevalent BYDV strain found in mainland Europe and the UK (BYDV-PAV) and several populations of the two most important vector species, *R. padi* (seven populations) and *S. avenae* (25 populations). Fig. 1 provides a graphical representation of our study system. Broadly, our results provide biological insights into the drivers behind variable transmission efficiency in an important vector-virus system.

**Fig. 1:**
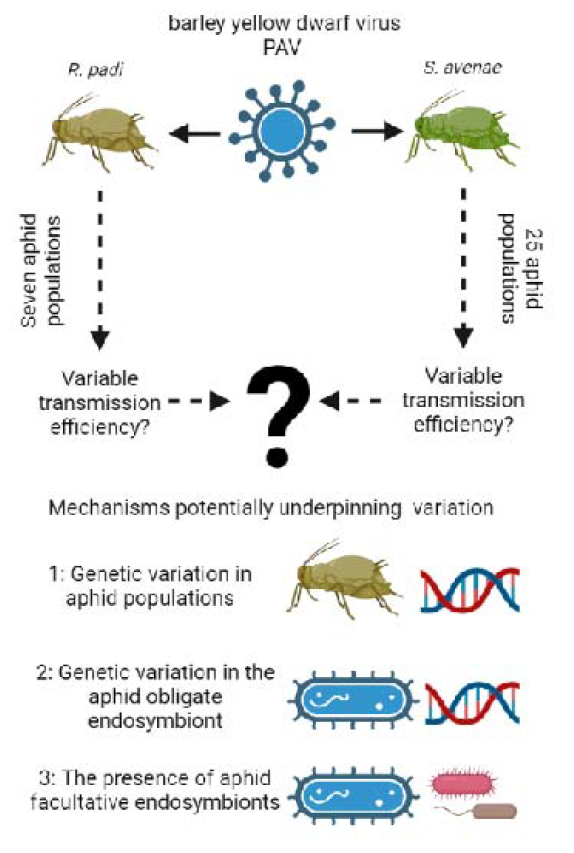
Graphical representation of the study system.

## Materials and Methods

### Rearing conditions and characterisation of aphid intra-species diversity

Aphid populations comprised 25 *S. avenae* populations and seven *R. padi* populations, an additional *R. padi* population (RP-12) was included in our genotyping analysis but was not included in the BYDV-PAV transmission studies. We retained this aphid in the study to increase the genetic data available when conducting our phylogenetic analyses. All aphid populations were maintained in cup cultures (similar to Leybourne, Bos [21]) under controlled environment conditions (18 ± 2 °C; 16:8 h L:D cycle) in a plant growth room on *Triticum aestivum* cv. Alcedo. The sampling location for all aphid populations, alongside their characterised facultative endosymbiont communities, was described previously [11]. The facultative endosymbiont associations are visualised in Fig. 2 (*R. padi*) and Fig. 3 (*S. avenae*).

**Fig. 2:**
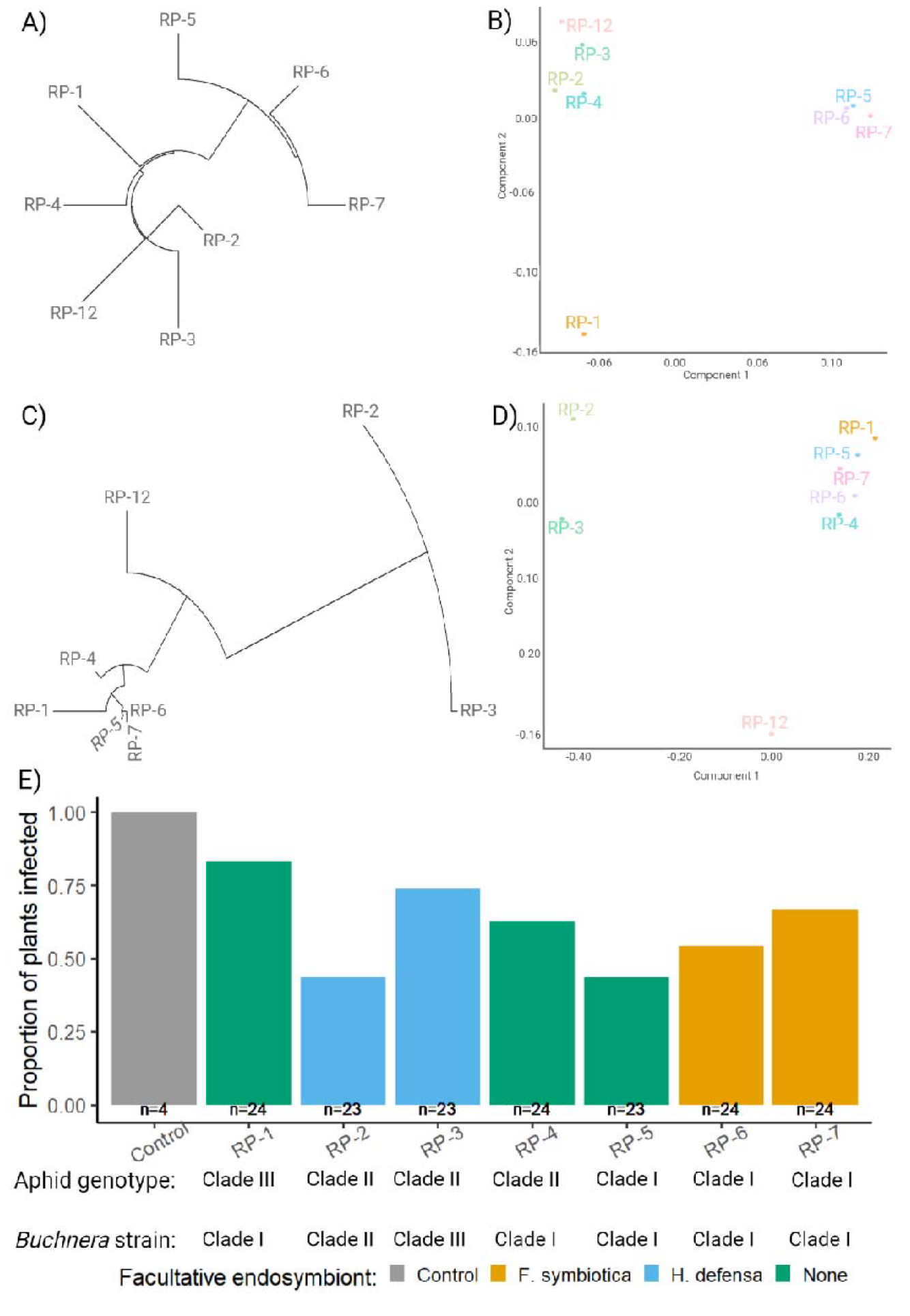
Newick tree (A, C) and MSDS clustering (B, D) for *R. padi* (A, B) and *B. aphidicola* (C, D) based on SNPs identified from Illumina sequencing. E shows the transmission efficiency (proportion of plants successfully infected with BYDV-PAV) for each *R. padi* population and the internal control; bar colour shows facultative endosymbiont presence and *n* represents the number of replicates.

**Fig. 3:**
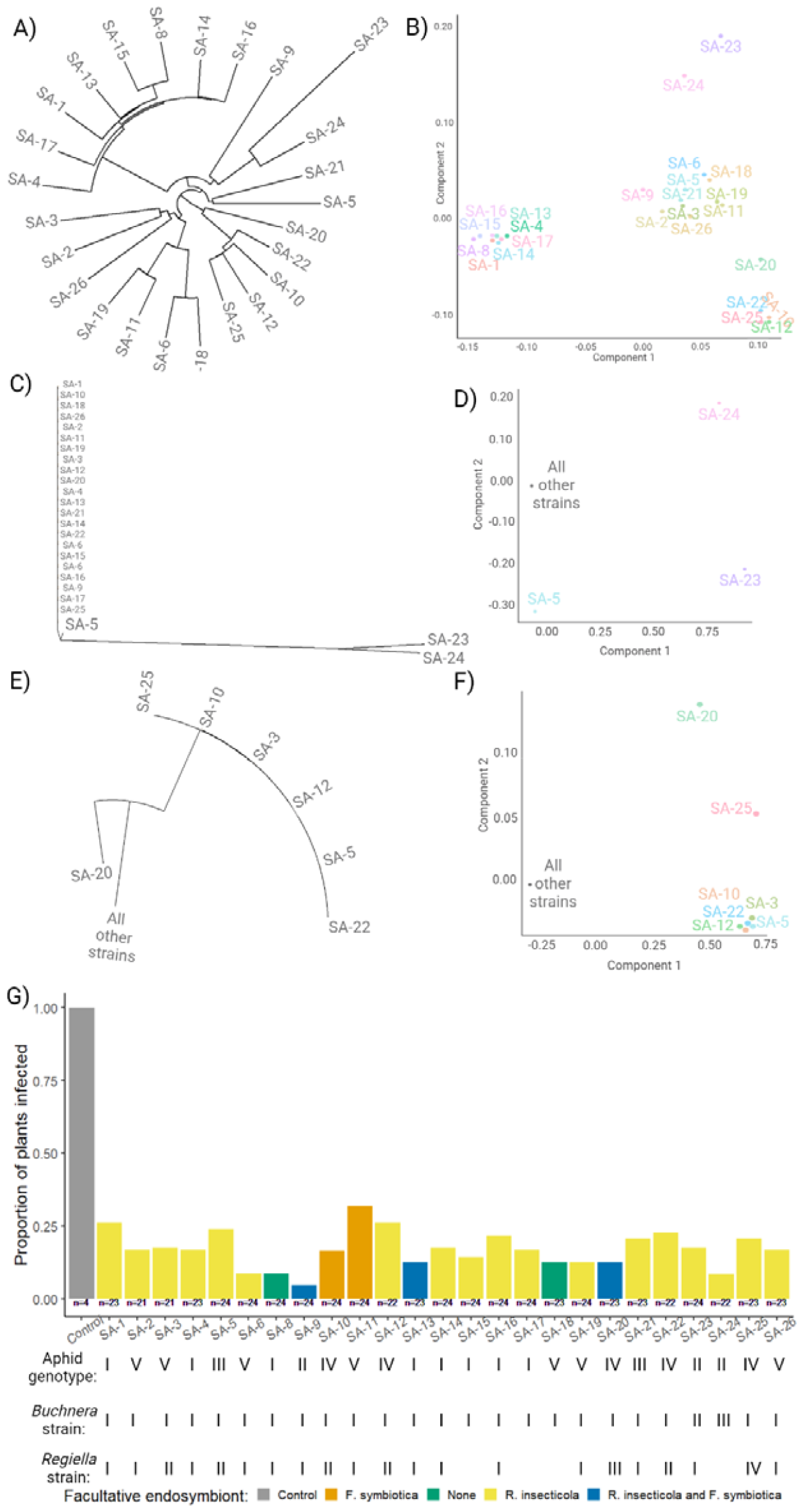
Newick tree (A, C, E) and MSDS clustering (B, D, F) for *S. avenae* (A, B), *B. aphidicola* (C, D), and *R. insecticola* (E, F) based on SNPs identified from Illumina sequencing. G shows the transmission efficiency (proportion of plants successfully infected with BYDV-PAV) for each *S. avenae* population and the internal control; bar colour shows facultative endosymbiont presence and *n* represents the number of replicates.

For aphid genotyping, approximately 40 aphids (mixed adults and nymphs) were collected into 96% molecular biology grade ethanol. Samples were sent to LGC Genomics GmbH (Berlin, Germany) for DNA extraction and sequencing (150 bp paired-end reads on an Illumina NextSeq 500/550 platform). All DNA extraction, library preparation, and sequencing were conducted by LGC Genomics GmbH. Data processing and SNP characterisation was carried out at the Centre for Genomics Research (The University of Liverpool).

### SNP calling

Aphid genomes were obtained from online databases: *S. avenae* genome (https://figshare.com/collections/Grain_aphid_Sitobion_avenae_genomics/5425896/1) [28], *R. padi* genome assembly 2.0 (https://bipaa.genouest.org/sp/rhopalosiphum_padi) [29]. Symbiont genomes were obtained from the NCBI database: *B. aphidicola* of *S. avenae* (accession: GCF_005082585.1), *B. aphidicola* of *R. padi* (accession: GCF_005080845.1), *F. symbiotica* (accession: GCF_003122425.1), *H. defensa* (accession: GCF_000021705.1), *R. insecticola* (accession: GCF_013373955.1). Aphid genomes were assessed for the presence of any symbiont genomes using blastn (using megablast algorithm [30], -evalue 1e-25) (v2.12.0+). *B. aphidicola* contigs were subsequently identified and removed from the *R. padi* assembly.

Reads were mapped with BWA MEM [31] (v0.7.17-r1188), and duplicate reads were marked with picard mark duplicates (v2.8.2). Variant calling and initial filtering were performed with samtools [32] (v1.6), bcftools [32] (v1.9), vcftools [33] (v0.1.16), and snpEFF [34] (v5). VCF’s were initially filtered with a minor allele frequency (MAF) of 0.05 to remove low-quality variants that are rare within the population, as well a low cut-off of depth 2. For phylogenetic tree inference, variants were retained where a genotype was called for each variant site in all samples. VCF’s were thinned using vcftools to 1000 or 5000 bp for symbiont or aphid genomes, respectively. Any resulting SNPs were retained for multidimensional scaling (MDS) plot analysis in plink [35] (v1.9), and Newick tree generation using the python package VCF kit [36] (v0.2.9).

We used plink to perform linear regression and assess for SNPs associated with BYDV transmission and BYDV titre. For aphid samples, instead of using thinned VCFs, variants were thinned using “--indep 50 5 2” to account for linkage disequilibrium (LD). LD pruning was not performed for symbiont samples. Phenotype association studies were performed in plink using the “--allow-no-sex --noweb --linear --ci 0.95” options. Variants with a P value < 0.05 were deemed to be of interest.

Phylogenetic distance between aphid and symbiont populations along the Newick tree and separation into distinct MDS clusters were used to assign our aphid populations into putative aphid genotype, the *B. aphidicola* and any associated secondary endosymbionts into putative microbial strains.

### BYDV-PAV transmission experiments

Apterous adult aphids from each population were randomly selected and placed onto BYDV-PAV infected *T. aestivum* cv. Alcedo plants and left to feed for 48 h; source plants had a mean relative virus titre of 2.97 ± 0.62 (measured by DAS-ELISA) and the BYDV-PAV culture held at the Julius Kühn Institut was used as a virus source [37]. Following this virus acquisition period, five adult apterous aphids from each population were removed and placed at the base of a wheat plant for 48 h, after which aphids were removed and plants were treated with insecticide; virus-carrying aphids from the BYDV-PAV stock culture were used as a control. The BYDV-susceptible wheat cultivar Alcedo was used in the BYDV transmission assays. Experimental plants were at the two-leaf stage (BBCH GS12) when challenged with BYDV-carrying aphids. Plants were retained in the controlled environment chamber for six weeks for virus incubation. After six weeks plants were screened for BYDV symptoms [5] and material was collected for a serological detection of BYDV infection via DAS-ELISA. The number of replicates per aphid population ranged from 21-24. Experiments were carried out in a controlled environment room (20 °C ± 2°C; 14:10 h L:D cycle).

For DAS-ELISA, 96-well polystyrene immunoassay microtiter plates were prepared by coating the plates with BYDV-specific polyclonal antibodies (IgG). BYDV-PAV IgG were prepared by the Julius Kühn Institut. The IgG concentration used was 1:200, diluted in ELISA coating buffer comprising: Na_2_CO_3_ (1.59 g / L), NaHCO_3_ (2.93 g / L), NaN_3_ (0.2 g / L); pH 9.6. 100 µL of IgG solution was added to each well, leaving two wells spare for blanks. Plates were incubated at 37 °C for 4 h in a moist chamber. After incubation, plates were emptied and washed four times with wash buffer (PBS-tween: 40 g NaCl, 7.2 g Na_2_HPO_4_-2H_2_O, 1.0 g KH_2_PO_4_, 1.0 g KCl in 5 L, with 2.5 mL Tween; pH 7.3) using a plate washer (Tecan Hydrospeed, Crailsheim, Germany). 50 mg of leaf tissue was sampled from each plant and placed in a 2 mL bead milling tube containing five steel beads. Samples were homogenised in 500 µL extraction buffer (wash buffer + 2% Polyvinylproline and 0.2% dry milk) through shaking in a Precellys® Evolution homogeniser for 30 s at 25,000 RPM. A 100 uL aliquot of homogenate was placed in an IgG-coated well, and three negative controls (uninfected plant tissue) and three positive controls were included in each plate. Plates were covered and incubated at 4-6 °C overnight.

After the overnight incubation plates were washed a further 5 times in wash buffer and the enzyme conjugate solution was added. The enzyme (alkaline phosphatase) conjugate solution comprised a 1:10,000 dilution of enzyme in extraction buffer. 100 uL of enzyme conjugate solution was added to each well and the plate was incubated at 37 °C for 4 h. Plates were washed a further four times and 200 µL Substrate buffer was added to each well. Substrate buffer comprised: 97 mL diethanolamine, 200 mg NaN_3_, and 203 mg MgCL_2_^*^6H_2_O in 1 L H_2_O + 1 mg / mL *p*-nitrophenyl phosphate (pNPP); pH 9.8. Plates were incubated in the dark for 60 minutes; endpoint extinction was measured at 405 nm using a plate reader (Tecan Sunrise). Two blank wells (containing substrate buffer only) were included per plate and all wells were blank corrected. The extinction intensity is a measure of the relative virus content/titer. The threshold for a positive BYDV-PAV infection was calculated at EXT of□>□0.06 (x(mean negative control)+3_∗_STD).

### Statistical analysis

All statistical analysis was carried out using R (v.4.3.0) [38] and R Studio (v.1.3.1093). The following additional packages were used to support data analysis and data visualisation: car v.3.0-11 [39], ggplot2 v.3.3.5 [40]. In all models, response variables included aphid genotype, *B. aphidicola* strain, and facultative endosymbiont presence; *B. aphidicola* strain and facultative endosymbiont presence were tested as nested variables within aphid genotype. In the *S. avenae* models *R. insecticola* strain was included as an additional explanatory variable (nested within aphid genotype).

BYDV transmission efficiency was analysed using general linear model fitted with a binomial distribution and a logit link. A binary value (1 = infected; 0 = uninfected) was modelled as the response variable and each aphid species was tested in a separate model. A Type II Wald χ2 analysis of deviance test was used to test the model. Differences in viral inoculation (ELISA titre) were examined using linear models. The BYDV titre of successfully-infected plants was modelled as the response variable and each aphid species was tested in a separate model. A Type II Anova was used to test the model.

## Results

### Genetic variation in R. padi influences BYDV transmission efficiency

From the Illumina data we identified 6,444 *R. padi* SNPs and used these to group the *R. padi* populations into three genetically similar clades (Fig. 2A, B). We used these clusters to call putative genotypes (Clades) for the *R. padi* populations. We also observed clustering for the obligatory endosymbiont *B. aphidicola* (Fig. 2C, D) and called putative strains for these based on 34 SNPs.

We detected significant variation in BYDV transmission efficiency between the *R. padi* populations examined (Fig. 2E), with differences attributed to aphid genotype (X^2^_2_ = 6.12; p = 0.046). On average, aphids in Clade I were the least efficient BYDV-PAV vectors and Clade III the most efficient vector. We identified 542 SNPs that potentially contribute to variable transmission efficiency in *R. padi. B. aphidciola* strain (X^2^_1_ = 5.53; p = 0.063) and facultative endosymbiont presence (X^2^_1_ = 2.38; p = 0.123) had no observable effect on BYDV transmission efficiency in *R. padi*. We did not detect any effect of aphid genotype (F_2.95_ = 1.92; p = 0.151), *B. aphidicola* strain (F_2.95_ = 0.61; p = 0.543) or facultative endosymbiont presence (F_1.95_ = 0.90; p = 0.345) on BYDV titre inoculated into the plant tissue following successful transmission (Fig. S1A).

### Transmission of BYDV-PAV by S. avenae is broadly inefficient and not affected by vector diversity

We identified 5,274 SNPs in our *S. avenae* Illumina data and the 25 *S. avenae* populations clustered into several clades (Fig. 3A, B). In contrast with *R. padi*, we did not observe a high level of genetic diversity across the obligatory endosymbiont *B. aphidicola*, with the majority of *B. aphidicola* strains grouping together based on eight SNPs (Fig. 3C, D). However, we detected genetic variation in the facultative endosymbiont, *R. insecticola*, with four putative strains called based on 29 SNPs (Fig. 3E, F).

For the *S. avenae* populations examined (Fig. 2G) we observed no effect of aphid genotype (X^2^_4_ = 5.71; p = 0.222), *B. aphidciola* strain (X^2^_2_ = 0.84; p = 0.656), facultative endosymbiont presence (X^2^_5_ = 1.74; p = 0.884), or *R. insecticola* strain (X^2^_4_ = 1.76; p = 0.778) on BYDV-PAV transmission efficiency. When compared with *R. padi* (Fig. 2E), all *S. avenae* clones examined (Fig. 3G) were more inefficient at vectoring BYDV-PAV. We did not detect any effect of aphid genotype (F_4.82_ = 0.85; p = 0.495), *B. aphidicola* strain (F_2,82_ = 0.30; p = 0.739), facultative endosymbiont presence (F_5,82_ = 0.86; p = 0.510), or *R. insecticola* strain (F_4.82_ = 0.67; p = 0.609) on BYDV titre inoculated into the plant tissue following a successful transmission (Fig. S1B).

## Discussion

Here we show that transmission efficiency of BYDV-PAV by *R. padi* is influenced by genetic variation within the vector population, and we identify 542 SNPs that are potentially involved in influencing BYDV-PAV transmission efficiency in *R. padi*. Our findings help disentangle the relationship between vector diversity and transmission efficiency of important plant viruses and provide insights that can guide future research endeavours.

A recent review synthesised information available on transmission efficiency of yellow dwarf virus species across the main cereal aphid vectors, including *R. padi* and *S. avenae* [5]. This synthesis identified significant variation in transmission efficiency across virus species and strains, vector species, and different clonal populations within a given vector species [5]. Variation in transmission efficiency in different vector-virus and virus-host (plant) combinations is unsurprising, as vector-virus relationships can be highly specific. Indeed, vectors are often characterised as efficient (competent, or compatible) or inefficient (incompetent, or incompatible) vectors for a given virus strain. For our study we selected *R. padi* and *S. avenae* as focal vector species as these species are considered to be important and efficient vectors for BYDV-PAV [5, 41, 42]. However, as we observed consistently low levels of transmission efficiency of BYDV-PAV by *S. avenae*, this aphid species might only be a moderately efficient vector for BYDV-PAV when compared with *R. padi*.

For *R. padi* we observed significant variation in BYDV-PAV transmission efficiency between the populations examined, with transmission efficiency ranging from *c*. 40-80%. This broadly supports previous observations that transmission efficiency can vary significantly between clonal populations for a given cereal aphid species [5, 37, 43]. Indeed, a recent synthesis showed that transmission efficiency of BYDV-PAV in *R. padi* can range from 0-100% (barley), 20-100% (oats), and 39-80% (wheat) [5]. Three mechanisms that potentially explain variations in transmission efficiency between populations within an aphid species were recently proposed [5]. These included: 1) Indirect effect of facultative endosymbionts through altered aphid feeding and probing behaviour; 2) Direct effect of *B. aphidicola* through variation in endosymbiont-derived chaperonin proteins; 3) Aphid genetic variation and the presence of vectoring alleles. Our study represents an examination of these hypotheses in *R. padi* and *S. avenae*, and identifies genetic diversity as a key factor underpinning transmission efficiency in *R. padi*. We excluded behavioural plasticity as a source for varying BYDV-PAV transmission efficiency, as it is low for feeding behaviour associated with virus acquisition and transmission in *S. avenae* [44].

Our observation of differential BYDV-PAV transmission efficiency across our *R. padi* genotypes complements results reported for the wheat aphid, *Schizaphis graminum* [27, 45-47]. Previous research used efficient and inefficient *Sc. graminum* populations and two other yellow dwarf virus species, CYDV-RPV (Solemoviridae : Polerovirus) and WYDV-SGV (Solemoviridae) to examine how genetic traits influence virus transmission efficiency [27, 45-48]. This process identified ‘vectoring alleles’ that underpin efficient transmission of CYDV-RPV in *Sc. graminum* [27] and provides evidence that genetic diversity within vector populations is a key driver of transmission efficiency [45-48], as also found here for our *R. padi* populations. We identified 542 SNPs in *R. padi* that are likely involved in underpinning the observed variation in BYDV-PAV transmission efficiency. However, it should be noted that our observations are based on a relatively small number of aphid populations, and that information on additional populations is required in order to fully elucidate the genetic traits underpinning transmission efficiency in *R. padi*. Nonetheless, to the best of our knowledge no other studies have characterised genetic diversity within different vector populations and linked this with variation in yellow dwarf virus transmission efficiency in *R. padi* [5]. Therefore, the previous insights gained in the *Sc. graminum* – CYDV-RPV and *Sc. graminum* – BYDV-SGV systems and our new observations in the *R. padi* – BYDV-PAV system represent important advancements in our understanding of the relationship between genetic diversity, vector-virus interactions, and transmission efficiency.

We found no evidence to support the two other hypotheses recently proposed [5]: 1) Indirect effect of facultative endosymbionts through altered aphid feeding and probing behaviour; 2) Direct effect of *B. aphidicola* through variation in endosymbiont-derived chaperonin proteins. However, some of the phenotypic traits conferred by facultative endosymbionts can act in a synergistic manner with host aphid genotype [22, 49]. Therefore, future work that explores endosymbiont effects on BYDV transmission in interaction with aphid genotype, for example through the elimination and introduction of endosymbionts via antimicrobial treatment and microinjection, while controlling for host aphid genotype, would enable a more robust examination of these hypotheses.

### Conclusion and future directions

Our work presents an investigation into the influence diversity in vector populations has on the transmission efficiency of an important cereal virus. We find that the two vector species examined, *R. padi* and *S. avenae*, can be broadly categorised into highly efficient and moderately efficient vectors of BYDV-PAV, respectively. In the efficient vector, *R. padi*, we identify significant variation in BYDV-PAV transmission efficiency and show that this is broadly driven by aphid genetic variation, with the population belonging to Clade III the more efficient vector. We identify 542 SNPs that are potentially involved in determining transmission efficiency in *R*. padi, although additional research that incorporates a greater number of *R. padi* populations is needed to confirm this. In the *R. padi* populations examined, we found no significant influence of *B. aphidicola* diversity or the presence of the facultative endosymbionts *H. defensa* and *F. symbiotica* on BYDV-PAV transmission. However, future work could disentangle potential interactive effects by investigating the potential aphid genotype x endosymbiont effects by manipulating the endosymbionts of aphids from Clade III (high efficiency) and Clade I (low efficiency) through elimination (antimicrobial treatment) and introduction (microinjection) of different *B. aphidicola* strains and facultative endosymbiont species.

## Acknowledgement

Illumina data analysis was carried out by the Centre for Genomic Research, which is based at the University of Liverpool. We would like to thank Kerstin Welzel (JKI) for rearing the aphid populations, and Evelyn Betke, Katharina A. Stein, and Heike Dobrowolski (JKI) for supporting the transmission experiments.

## Data availability

Illumina data have been deposited to the European Nucleotide Archive (ENA) database, project ID PRJEB72361. Informatics code used in the project are available via GitHub (https://github.com/hlmwhite/BYDV_code). Transmission data and code are available via the University of Liverpool’s Data Catalogue (doi: 10.17638/datacat.liverpool.ac.uk/2607).

