## Supplementary material for "Genetic diversity in vector populations influences the transmission efficiency of an important plant virus": Fig. S1

**Fig. S1:** ELISA titre for BYDV-infected plants following challenge with BYDV-carrying *R. padi* (A) and *S. avenae* (B). Plot colour shows facultative endosymbiont presence.

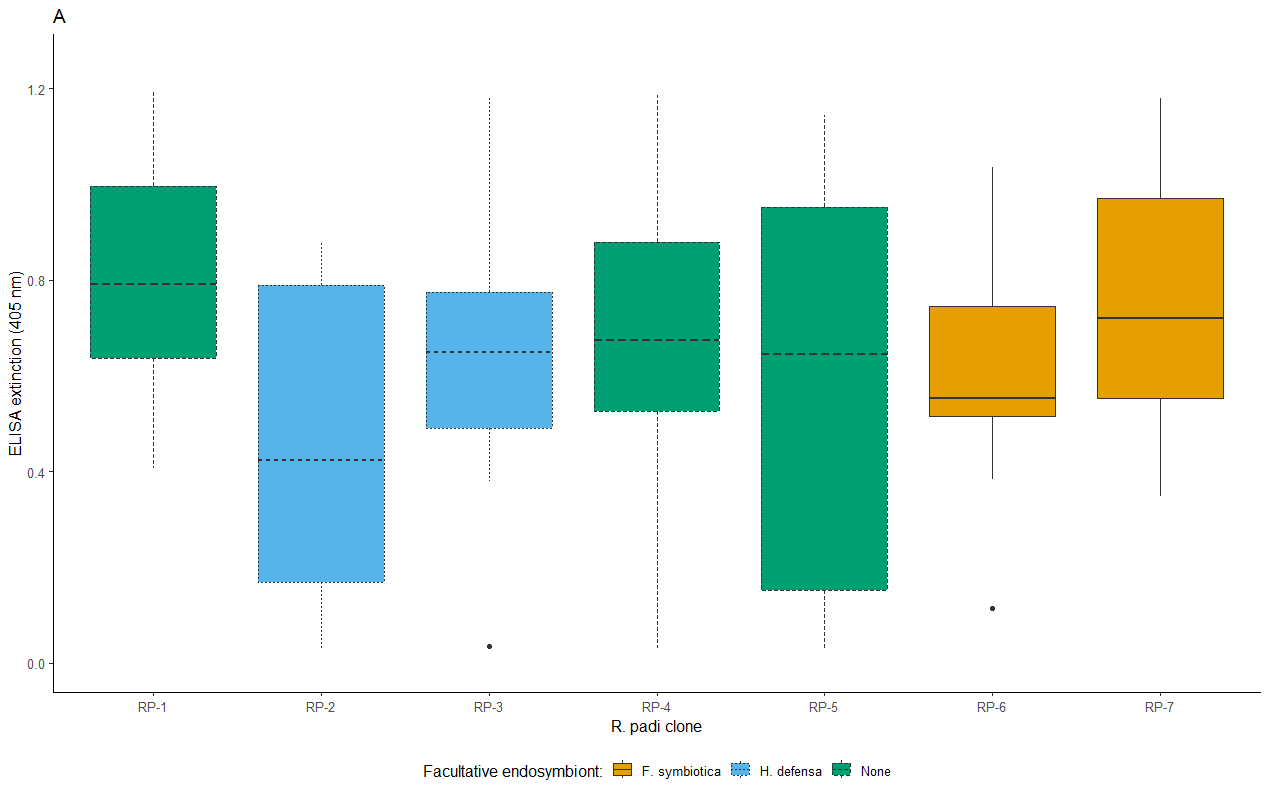

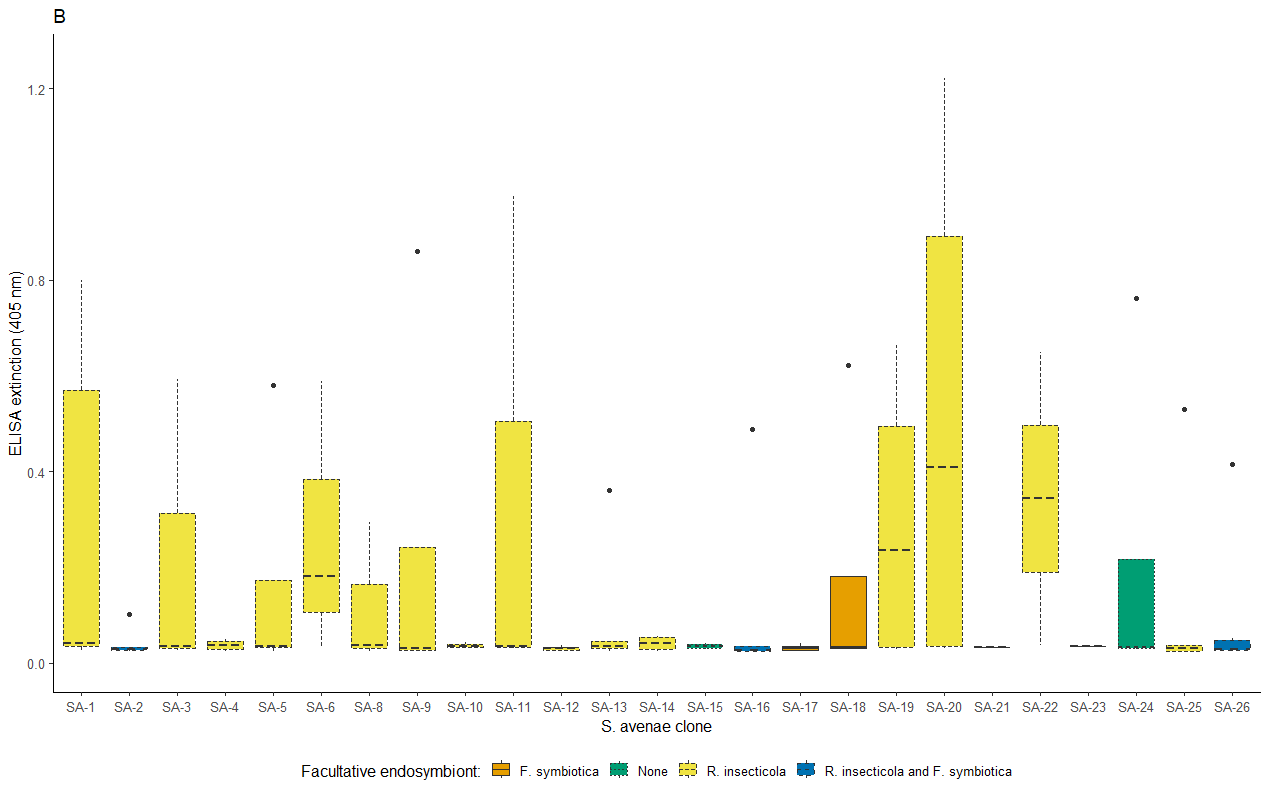
